# Additional layer of regulation via convergent gene orientation in yeasts

**DOI:** 10.1101/551689

**Authors:** Jules Gilet, Romain Conte, Claire Torchet, Lionel Benard, Ingrid Lafontaine

## Abstract

Convergent gene pairs can produce transcripts with complementary sequences. We had shown that mRNA duplexes form *in vivo* in *Saccharomyces cerevisiae* via interactions of mRNA overlapping 3’-ends and can lead to post-transcriptional regulatory events. Here we show that mRNA duplex formation is restricted to convergent genes separated by short intergenic distance, independently of their 3’-UTR length. We disclose an enrichment in genes involved in biological processes related to stress among these convergent genes. They are markedly conserved in convergent orientation in budding yeasts, meaning that this mode of post-transcriptional regulation could be shared in these organisms, conferring an additional level for modulating stress response. We thus investigated the mechanistic advantages potentially conferred by 3’-UTR mRNA interactions. Analysis of genome-wide transcriptome data revealed that Pat1 and Lsm1 factors, having 3’-UTR binding preference and participating to the remodeling of messenger ribonucleoprotein particles, bind differently these messenger interacting mRNAs (mimRNAs) forming duplexes in comparison to mRNAs that do not interact (solo mRNAs). Functionally, mimRNAs show limited translational repression upon stress. We thus propose that mRNA duplex formation modulates the regulation of mRNA expression by limiting their access to translational repressors. Our results thus show that post-transcriptional regulation is an additional factor that determines the order of coding genes.

## Introduction

The transcriptional orientation of genes relative to their adjacent gene neighbors along the chromosome can be either co-orientation (→→), divergence (←→) or convergence (→←) (fig. 1A). This genomic neighborhood may reveal functional constraints. In eukaryotes, neighboring genes are likely to be co-expressed, independently of their relative orientation (Cohen et al. 2000; Hurst et al. 2004; Michalak 2008). To date, most attention has been devoted to the link between genomic neighborhood and co-transcriptional regulation. Co-orientation can allow co-regulation of transcription of the two genes by a single promoter in an operon-like fashion (Osbourn and Field 2009) and divergence can allow co-regulation of transcription by means of a bi-directional promoter (Wei et al. 2011). In the case of convergent gene pairs that do not share any promoter region, co-transcription could be mediated by chromatin effects rather than by direct interactions (Chen et al. 2010). Most importantly, there is increasing evidence that convergent gene orientation can also mediate regulation at the post-transcriptional level. Transcriptome analyses have shown that convergent gene pairs can produce tail-to-tail 3’-overlapping mRNA pairs that can theoretically form mRNA duplexes in *S. cerevisiae* (Pelechano and Steinmetz 2013; Wilkening et al. 2013) and in other eukaryotes (Jen et al. 2005; Makalowska et al. 2005; Sanna et al. 2008). We previously demonstrated that such mRNAs duplexes exist extensively and can interact in the cytoplasm in *Saccharomyces cerevisiae* (Sinturel et al. 2015). Some of these mRNA-mRNA interactions are apparently strong enough to promote post-transcriptional regulatory events by blocking ribosome elongation, as illustrated by mRNA duplex formation between OCA2 and POR1 mimRNAs. Indeed, OCA2 mRNA was previously shown to overlap the protein coding sequence of POR1 mRNA, block ribosome elongation and trigger the no-go decay pathway (Doma and Parker 2006; Passos et al. 2009; Sinturel et al. 2015).

**Figure 1.**
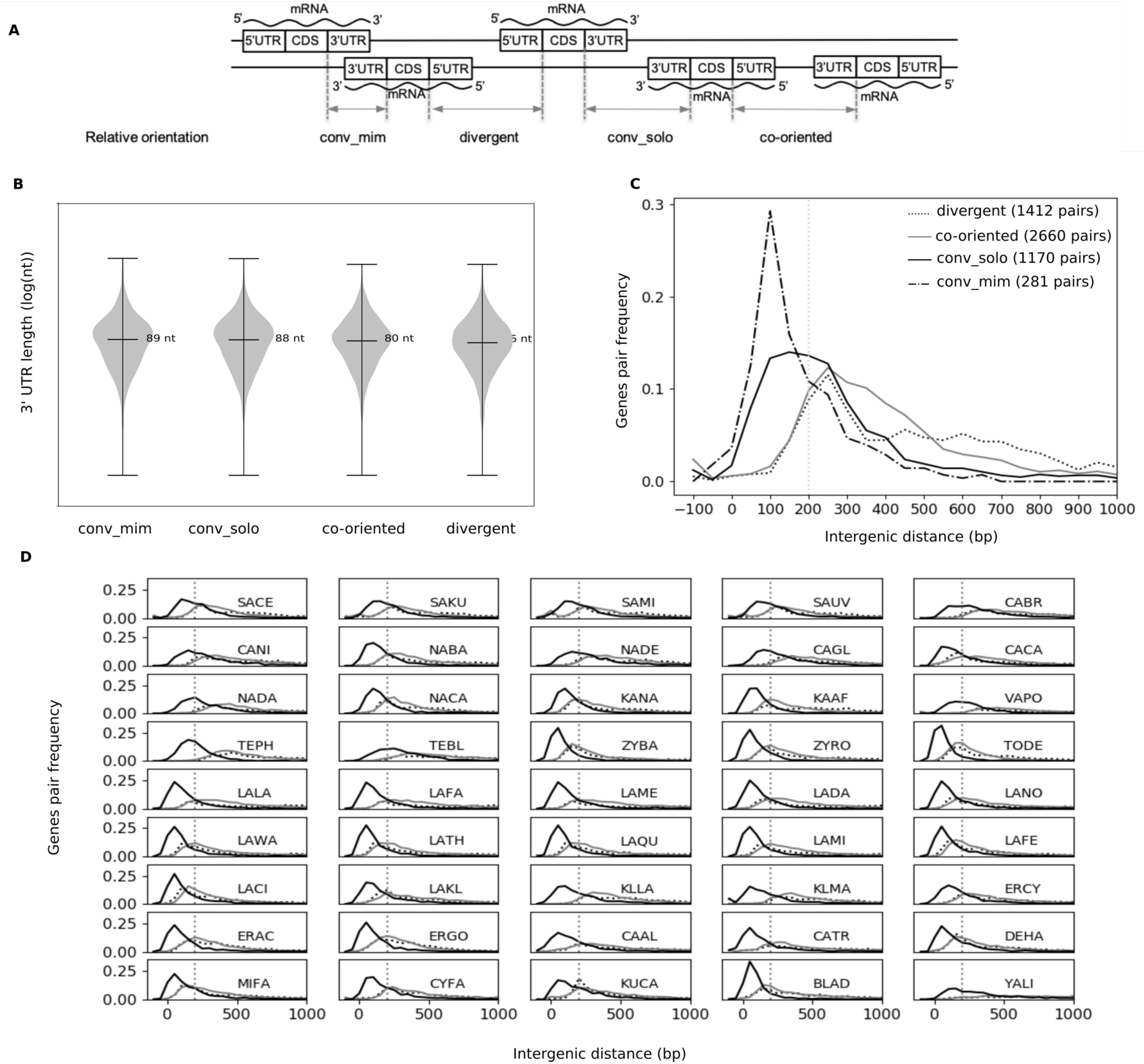
mRNA duplexes form at small intergenic distances, independently of their lengths. (A) Schematic representation of relative orientation of adjacent genes. Intergenes are delimited by dashed blue lines and their distances indicated by double arrows. *conv_mim*: convergent gene pairs producing mimRNAs forming experimentally validated mRNA duplexes, *conv_solo*: convergent gene pairs with no experimentally validated RNA duplexes, *divergent*: genes in divergent orientation. *co-oriented*: genes in co-orientation. (B) mRNA 3’-UTR lengths (logarithmic scale) for different gene groups in *S. cerevisiae* taken from (Nagalakshmi et al. 2008). Median values are indicated for each group. (C) Gene pair frequency distribution in function of their intergenic distance for conv_mim (dot-tick black line), conv_solo (black line) co-oriented (grey line) and divergent (black dotted line) gene pairs in *S. cerevisiae*. (D) Gene pair frequency distribution in function of the intergenic distance for convergent pairs (black line), co-oriented pairs (grey line) and divergent pairs (black dotted line) in the 45 species studied. Species are named with a 4-letter code available in supplementary table S2 and ordered according to their evolutionary distance from *S. cerevisiae*. The dashed horizontal line indicates 200 bp, the distance below which a majority of conv_mim are found, in contrast to conv_solo in *S. cerevisiae*.

We thus hypothesize that mRNA duplex formation can modulate interactions with RNA binding proteins that preferentially bind at mRNA 3’-end, like Pat1 and Lsm1. Pat1 and Lsm1 are considered as main translation repressors activated during stress and also as key players in 5’ to 3’ mRNA decay, linking deadenylation to decapping (Tharun and Parker 2001; Tharun et al. 2000; Chowdhury and Tharun 2009). Pat1 and Lsm1 are components of messenger ribonucleoprotein particles (mRNP), named P-bodies, which contain translational repressors and the mRNA decay machinery (Mitchell and Parker 2014).

If genomic neighborhood plays some critical role in gene expression, these should be conserved when under selection and detected as conservation of microsynteny. In addition, favorable new genomic neighborhoods, obtained by chromosomal rearrangements, should also appear and be selected for during evolution. Although there is evidence for cis-regulatory constraints on gene order, our understanding of the determinants of the evolution of gene order in eukaryotes is still limited. Globally, gene pair conservation decreases as intergenic distance increases (Hurst et al. 2002; Poyatos and Hurst 2007). In yeasts, gene pairs that are highly co-expressed are more conserved than gene pairs that are not co-expressed and it has been reported that only divergent gene pairs are under selection for high co-expression (Kensche et al. 2008; Wang et al. 2011a; Yan et al. 2016). However, the co-expression of linked genes persists long after their separation by chromosomal rearrangements whatever their original relative orientation, and natural selection often favors chromosomal rearrangements in which co-expressed genes become neighbors. Thus, selectively favorable co-expression appears not to be restricted to bi-directional promoters (Wang et al. 2011a).

In order to determine the possible role of mRNA duplex formation in gene regulation, we performed a genomic analysis of genes producing mimRNAs on an evolutionary perspective to determine i) the extent to which mRNA duplexes could form in 45 budding yeasts (Saccharomycotina subphylum), covering an evolutionary distance of *ca.* 300 MYA (Marcet-Houben and Gabaldón 2015), ii) the functional properties of genes producing mimRNAs by a Gene Ontology (GO) enrichment analysis, iii) the conservation of convergent orientation among yeasts as a proxy for their functional significance. We also compared the properties of mimRNAs and solo mRNAs that do not form duplexes. First by analyzing cross-linking immunoprecipitation (CLIP) data used to map the interaction sites of Pat1 and Lsm1 on mRNAs (Mitchell et al. 2013) in condition of glucose deprivation, a condition triggering the formations of P-bodies, particularly requiring Pat1 and Lsm1. Secondly by analyzing ribosome loading data used to determine the ability of Pat1 and Lsm1 factors to repress translation of mRNAs in stress conditions (Garre et al. 2018).

Our results show that mRNA duplexes form between mimRNAs of genes that are less than 200 bp apart, independently of the length of their 3’-UTR. They are functionally enriched in biological processes occurring during response to stress in *S. cerevisiae* and provided orthology-function relationships are preserved, it is also the case in many yeasts for convergent genes less than 200 bp, theoretically able to form mRNA duplexes. We propose that mRNA– mRNA interactions can interfere with solo mRNP remodelers such as Pat1 and Lsm1, contributing to limit the translational repression on mRNA duplexes and thus participating in modulating gene expression upon stress. Furthermore, convergent orientation between neighboring genes is in general more conserved at short intergenic distances than co-oriented or divergent orientation in all 45 studied genomes, which suggests that convergent orientation allowing post-translation regulation of mRNA of genes involved in stress response is widely shared.

## Results

### RNA duplexes in *S. cerevisiae* occur between genes separated by short intergenic distances

In *S. cerevisiae*, 281 pairs of mRNA duplexes have been determined experimentally (see Materials and Methods, supplementary table S1 and Sinturel et al. 2015). The distribution of 3’-UTR length of the 562 corresponding mimRNAs is not statistically different from the distributions of the other solo mRNAs (Mann-Whitney tests, p-value > 0.05, fig. 1B). Conversely, the intergenic distances, defined as the distance between CDS regions of adjacent genes (fig. 1A), are shorter between convergent genes producing mimRNAs (median 155 bp) than between convergent genes producing solo mRNAs (median 236 bp) (Mann Whitney test, p-value <10^-13^, fig. 1C). We thus chose 200 bp as an appropriate distance cut-off for which a majority of experimentally validated mimRNAs are found (63.2%) in contrast with convergent genes producing solo mRNAs which are only 40.3% (supplementary fig. S1A). Note that 18% of the convergent gene pairs with intergenic distances below 200 bp correspond to validated mimRNAs and only 7% correspond to validate solo mRNAs. This suggests that the short intergenic distances, between convergent genes are the major determinant for mRNA duplex formation. We note that the bimodal distribution of intergenic lengths among divergent gene pairs is in agreement with previous reports (Hermsen et al. 2008).

The reconstructed phylogenetic tree of the 45 yeasts we studied, congruent with the backbone of the Saccharomycotina phylogeny (Shen et al. 2016), is presented in fig. 2. Within these genomes, convergent gene pairs are separated by the smallest intergenic distances, with a median of 158 bp, compared to co-oriented gene pairs (median of 405 bp) or divergent gene pairs (median of 517 bp) (supplementary table S2), a trend previously observed (Chen et al. 2011). Within 31 out of the 45 studied genomes, the 51st percentile of the intergenic distances between convergent genes is below 200 bp. In addition, a comparative transcriptomic analysis in different yeasts revealed that 3’-UTR lengths are also broadly similar (Moqtaderi et al. 2013), demonstrating that the majority of convergent transcripts overlap and are theoretically able to form mRNA duplexes. In 36 out of 45 genomes, the proportion of co-oriented gene pairs are significantly smaller than expected under a neutral model of gene order evolution, where genes would be equally distributed among the two DNA strands (50% of co-oriented, 25% of divergent and convergent) (supplementary table S3). Neighboring genes are then more often encoded on opposite strands, probably due to a greater impact of bidirectional promoters and of chromatin context for transcriptional regulation, or a greater impact of mRNA duplex formation for post-transcriptional regulation. Interestingly, in 27 of these 36 genomes, the proportion of convergent pairs is higher than those of divergent pairs.

**Figure 2.**
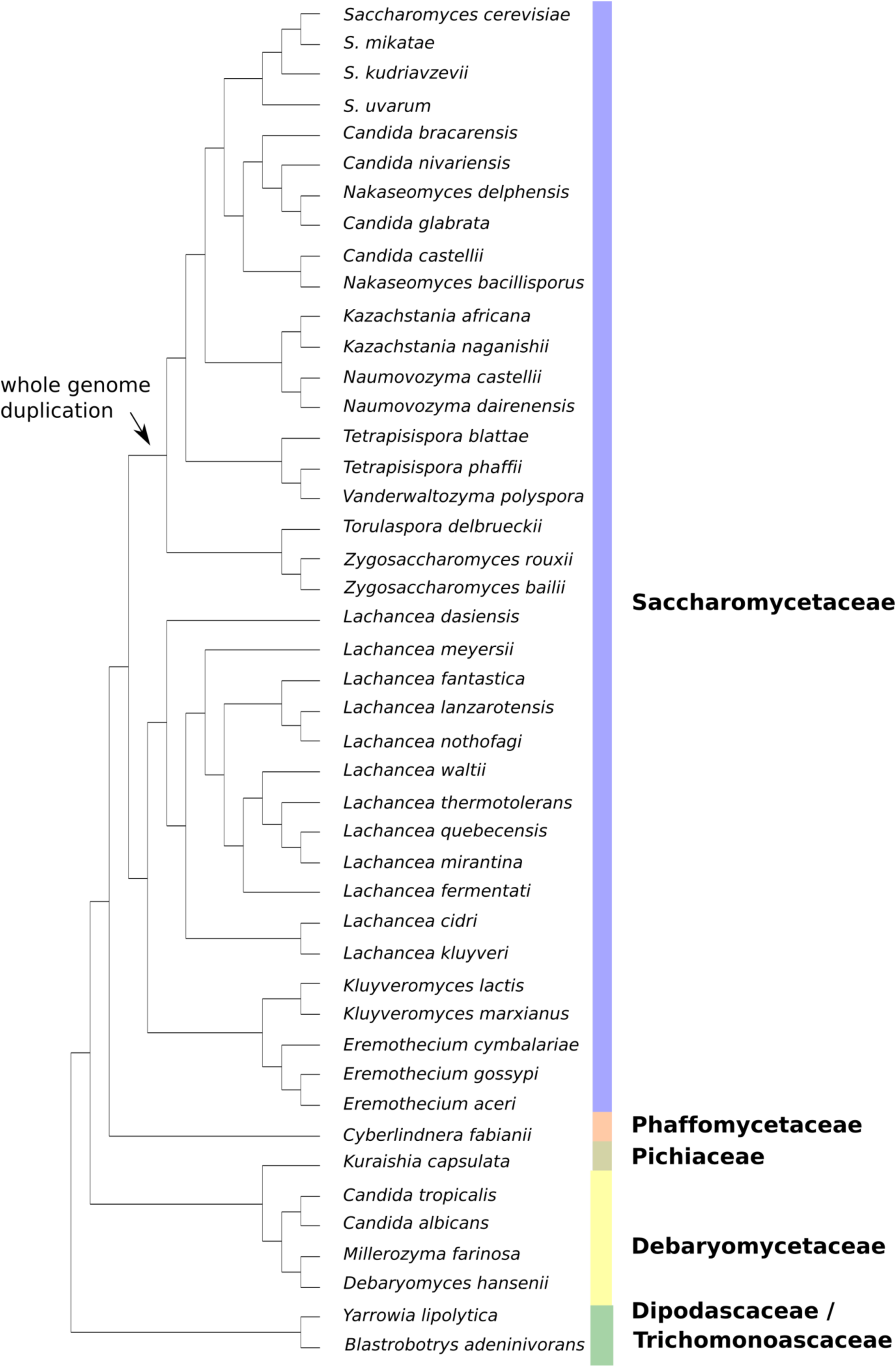
Phylogenetic relationships of the 45 Saccharomycotina yeasts species studied. Phylogeny of 45 Saccharomycotina species inferred from a maximum likelihood analysis based on the concatenated alignment of 224 groups of syntenic homologs present in every genome (See Materials and Methods).

### GO analysis of convergent genes

According to the Yeast GO Slim annotation (see Materials and Methods), the 365 genes producing experimentally validated mimRNAs in *Saccharomyces cerevisiae* are significantly enriched (more than 2-fold enrichment, hypergeometric test, adjusted p-values < 0.01) into cellular response to DNA damage stimulus, DNA metabolism (repair, recombination and replication) together with mRNA processing and RNA splicing (Table 1). We found the same results by estimating the probability of occurrence of each GO Slim term among 365 randomly selected genes (permutation p-values < 0.01) (supplementary table S4). Among the 919 convergent genes less than 200 bp apart in *Saccharomyces cerevisiae*, which are theoretically able to produce mimRNAs but not experimentally validated in (Sinturel et al. 2015), there is also an enrichment for DNA repair and cellular response to DNA damage stimulus (permutation p-value < 0.05) (supplementary table S4). On the contrary, there is no such enrichment among genes producing solo mRNAs whether being i) the 248 convergent genes that were experimentally validated as producing only solo mRNAs, ii) the convergent genes more than 200 bp apart and not experimentally validated and iii) the genes in co-oriented and divergent relative orientation that are less than 200 bp apart (supplementary table S4).

**Table 1.** GO Slim terms of biological processes significantly enriched (p-value < 0.01) more than 2-fold in *S. cerevisiae* mimRNAs validated experimentally.

| GO Slim term | # all genes<sup>a</sup> | # conv_<br>mimRNA<sup>b</sup> | fold<br>enrich.<sup>c</sup> | adj. p-value<sup>d</sup> |
| --- | --- | --- | --- | --- |
| DNA repair | 245 | 32 | 2.4 | 3.03E-04 |
| Cellular response to DNA damage stimulus | 299 | 33 | 2 | 2.12E-03 |
| DNA replication | 152 | 18 | 2.2 | 8.73E-03 |
| DNA recombination | 174 | 20 | 2.1 | 8.73E-03 |
| mRNA processing | 167 | 19 | 2.1 | 8.73E-03 |
| RNA splicing | 134 | 17 | 2.3 | 8.73E-03 |

Note - <sup>a</sup> number of GO slim terms in all *S. cerevisiae* genes (total number of annotated genes: 6320). <sup>b</sup> number of GO slim terms in *S. cerevisiae* genes producing mimRNAs (total number of annotated genes producing mimRNAs: 341). <sup>c</sup> fold-enrichment (frequency ratio in conv\_mimRNA and in all genes). <sup>d</sup> p-value of the hypergeometric test adjusted with the Benjamini-Hochberg procedure for multiple testing (see Materials and Methods).

For each of the other Saccharomycotina species, we determined the functional distribution of GO Slim terms among the total number of convergent genes with an identified ortholog in *S. cerevisiae* (N_conv_) -assuming they share the same function as their *S. cerevisiae* ortholog-that are less than 200 bp apart and thus theoretically able to produce mimRNAs that form mRNA duplexes (see previous section). We next calculated the probability of occurrence of each GO Slim term among N_conv_ randomly selected genes with an identified *S. cerevisiae* ortholog (fig. 3 and supplementary table S5). As is the case in *S. cerevisiae*, there is a significant enrichment (permutation p-value < 0.05) for cellular response to DNA damage stimulus, DNA repair and RNA splicing in at least 18 other yeast genomes. In addition, mRNA modification, tRNA processing, chromosome segregation and protein complex biogenesis are also enriched in more than one third of the yeast species. Such functional enrichment of terms that could be linked to stress response among convergent genes theoretically able to form RNA duplexes indicates that their mode of post-transcriptional regulation could be shared in most of these species.

**Figure 3.**
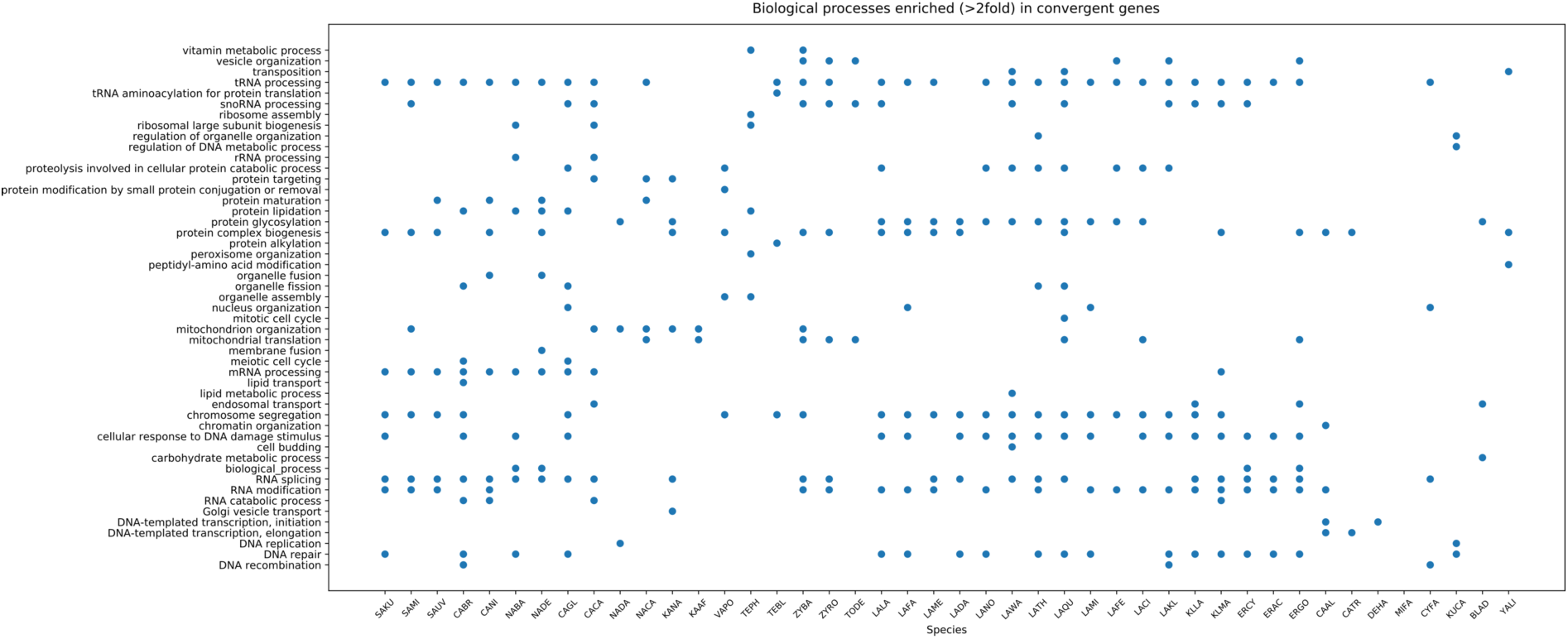
Functional enrichment in convergent genes. Biological process GO Slim terms with a significant enrichment (> 2 fold with a p-value < 0.05) among convergent genes compared to the whole gene population. *S. cerevisiae* GO slim terms have been attributed to orthologs in the 44 other species. Species are named with a 4-letter code available in supplementary table S2 and ordered according to their evolutionary distance from *S. cerevisiae*.

### Convergent relative orientation is more conserved than divergent and co-oriented relative orientations at short intergenic distances

A further insight at the potential importance of convergent orientation is its conservation during evolution. We defined the orthologs between each pair of the 45 yeast genomes (see Materials and Methods), and determined their relative orientation to their gene neighbors in the two considered species.

We first considered a conserved gene orientation when two orthologs share the same relative orientation with respect to their 3’ neighbor in both genomes, independently of the orthology relationship of the neighboring genes (genomic context). It allows one to estimate the extent at which the relative orientation is functionally important in itself, whether it being acquired from the ancestor or by chromosomal rearrangements. On average, without considering intergenic distances, co-orientation is the most conserved gene orientation (78%) followed by convergence (75%) and divergence (72%) (supplementary table S6). In all cases, conservation decreases as the evolutionary distance between species increases (fig. 4A, left panel). We then estimated the expected conservation levels under a null model by randomly assigning orthologous pairs between each species in 500 simulation rounds (fig. 4A, right panel). These data clearly show that the observed levels are higher than expected by chance. At four large pairwise evolutionary distances (1, 1.13 and 1.5, see supplementary Table S9) co-orientation is the less conserved orientation and convergence the most conserved one (fig. 4A, left). These distances always involve the species that are the most isolated from all the other studied species: *Cyberlindnera fabianii,* the only species from the *Phaffomycetaceae* clade studied (Kurtzman et al. 2008), and *Y. Lipolytica* and *B. adeninivorans* belonging to the most distant clade from all the others (fig. 2). These three species also share the lowest numbers of ortholog with all other species: 804, 1273 and 1722 orthologs shared with another species for *Y. Lipolytican, A. adeninivorans* and *C. fabiani* respectively, for an average of 2874 for the 45 species studied. This particular trend could reflect either a sampling bias or that the most conserved gene orientation between different yeast clades is the convergent one. For microsynteny, convergent orientation is the most conserved one at each evolutionary distance, which is also expected to a much lower extent under the null model (fig. 4B).

**Figure 4.**
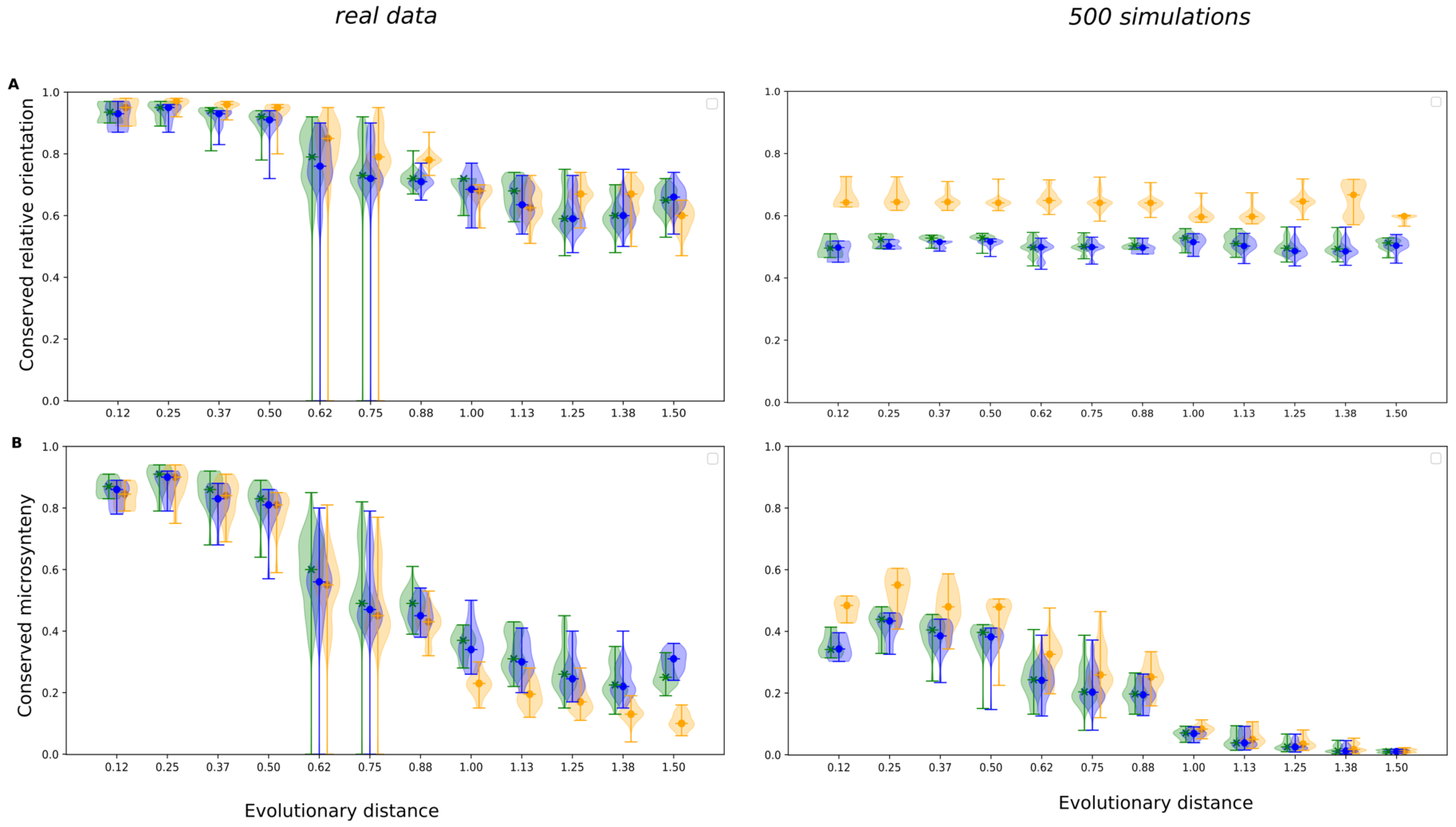
The conservation of relative orientation of orthologs decreases with the evolutionary distance between the species. (A) Frequency distributions of conserved relative orientation between two species as a function of their evolutionary distance (supplementary Table S9). (B) Frequency distribution of conserved microsynteny between two species as a function of their evolutionary distance. Green: genes in convergent orientation (conv); orange: co-oriented genes (coor) and blue: divergent genes (div). Left panels: distributions in the real genomes. Right panels: distributions in the 500 simulations experiment in which orthology relationships are at random (see Materials and Methods). See supplementary table S6 for the total number of conserved pairwise orthologs over the 990 pairs of species at each evolutionary range. Observed distributions of convergent pairs are different compared to distributions of co-oriented and divergent pairs at any given evolutionary interval (Mann-Whitney tests, p-value < 10^-3^).

As convergent genes are separated by smaller intergenic regions than divergent and co-oriented genes, we looked whether the physical proximity of convergent genes was the main explanation to their preferential conservation. To that end, we estimated the proportion of conserved gene relative orientation in windows of non-overlapping intergenic distances ranging from 0 to 1000 bp (supplementary table S10). Strikingly, for intergenic distances lower than 200 bp, convergent gene orientation, thus theoretically able to produce mimRNAs that form mRNA duplexes, is more conserved than divergent and co-oriented gene orientations (Mann-Whitney tests p-values < 10^-200^) (fig. 5A, panel 1 and 3). This higher conservation of convergent gene pairs is not due to their lower intergenic distances compared to co-oriented and divergent pairs (average distance of 120, 142 and 147 respectively) as it holds also for intergenic distances between 144 and 200 bp (Mann-Whitney tests p-values < 10^-200^) (fig. 5C, left). Above 200 bp, there is an opposite trend, the convergent gene orientation being less conserved (Mann-Whitney tests p-values < 10^-6^). It is of note that the above observations hold when considering species pairs that diverged before or after the whole genome duplication event that occurred during yeast evolution (pre-WGD species or post-WGD species) and is much higher than expected by chance alone (fig. 5A). Similar observations are made when grouping species pairs by evolutionary distances, thus reducing the bias that can be introduced when considering extant species pairs (supplementary fig. S2).

**Figure 5.**
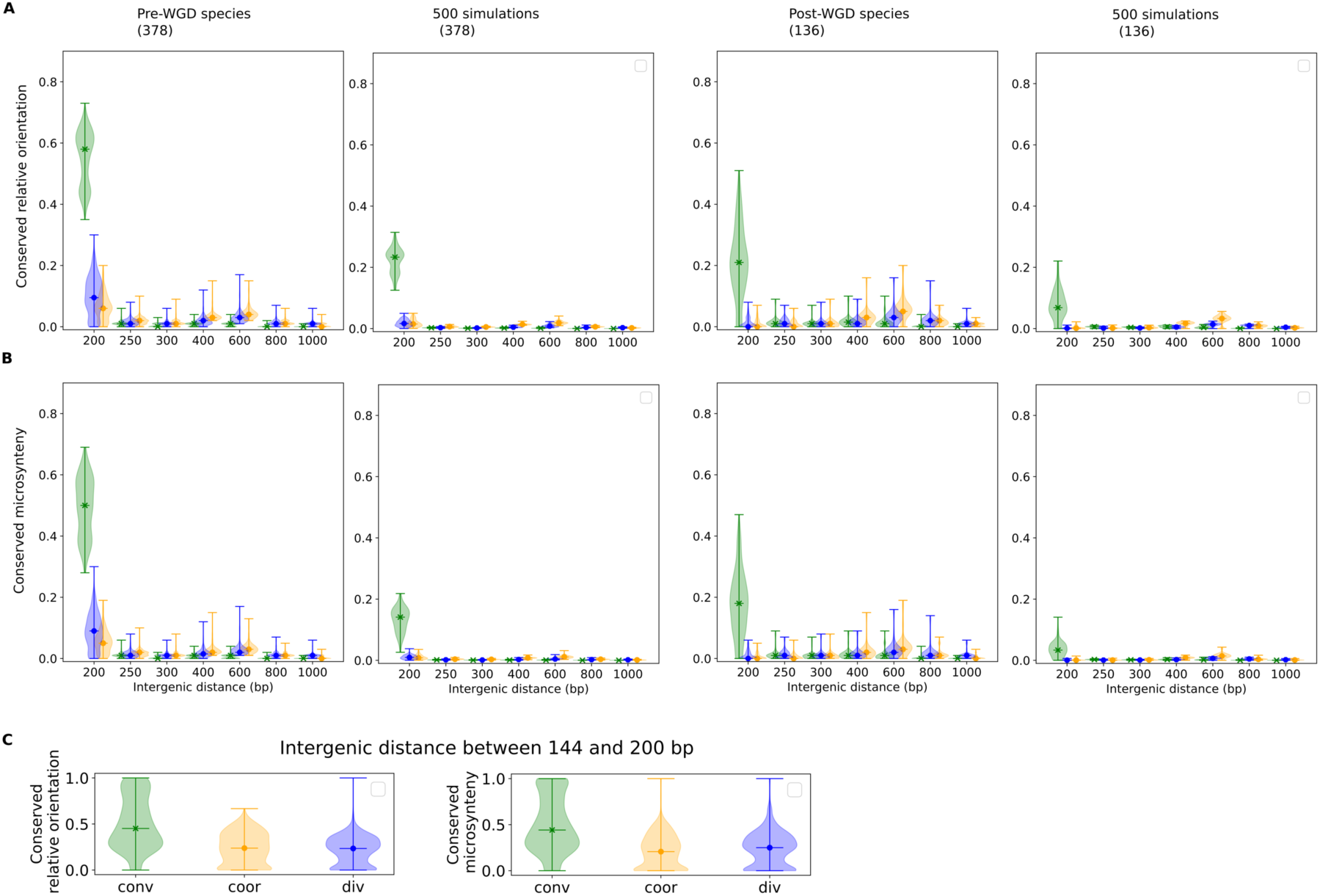
The conservation of relative orientation is higher for convergent pairs less than 200 bp apart. (A) Frequency distribution of conserved relative orientation in function of the length of the intergenic distance separating the genes in a pair. (B) Frequency distribution of conserved microsynteny as a function of the length of the intergenic distance separating the genes in a pair. Same color legend as in fig. 4. Distributions for species pairs that diverge before and after the whole genome duplication event (pre-WGD and post-WGD) are given in first and third panels, respectively. Distributions for the 500 simulations experiment for pre-WGD and post-WGD species pairs are given in second and fourth panel, respectively. Number of species pairs are given in parenthesis above panels. For each intergenic distance interval, the distribution for the convergent pairs is significantly different from the one of co-oriented and divergent pairs (Mann-Whitney tests, p-value < 0.05). See supplementary tables S7 and S8 for numbers of conserved orthologs at each intergenic distance intervals. (C) Frequency distribution of conserved relative orientation (left) and conserved microsynteny (left) of gene pairs separated by intergenic distances ranging from 144 to 200 bp, among orthologs conserved in same relative orientation/in microsynteny in this interval. The distribution of convergent pairs are different compared to distributions of co-oriented and divergent pairs (Mann-Whitney tests, p-value < 10^-200^).

We next considered a conserved microsynteny, when two orthologs share the same relative orientation with respect to their 3’ neighbor in both genomes, and when the neighboring genes are also orthologs, which corresponds to the conservation of the genomic context from their common ancestor only, see Materials and Methods. The conservation of microsynteny for convergent orientation remains the highest (fig. 5B) but to a lower extent than in the case of conservation of the genomic context (fig. 5A), for both pre-WGD and post-WGD species pairs. This holds as well between pairs of species at similar evolutionary distance (supplementary fig. S3). This holds also for intergenic distances between 144 and 200 bp (Mann-Whitney tests p-values < 10^-200^) (fig. 5C, right). As expected under a neutral model of evolution of gene order, the probability of gene pair conservation decreases with the length of the intergenic region between the genes for all types of pairs because the probability of recombination between two genes increases with their spacing. It has been shown indeed that intergenic distance is the major determinant of gene pairs conservation in yeasts (Poyatos and Hurst 2007). However, at small intergenic distances, the convergent pairs are less prone to recombination than co-oriented and divergent ones. This either reflects a selective pressure to maintain convergent pairs at small intergenic distances allowing RNA duplexes formation, and/or a counterselection of co-oriented and divergent pairs.

In summary, at small intergenic distances which allow for mRNA duplex formation, the conservation of microsynteny in convergent orientation appears not neutral, and the conservation of the genomic context in convergent orientation is even more important, suggesting that convergent gene pairs are both conserved and recruited by chromosomal rearrangements for functional constraints. At such intergenic distances, genes producing validated solo mRNAs show no conservation, in contrast to genes producing validated mimRNAs (supplementary fig. S1A and B), which argues for mRNA duplex formation being not only determined by small intergenic distances, but also by a selection pressure on a subset of genes found in this configuration.

### mRNA duplexes limit Lsm1 and Pat1 interactions in 3’-UTRs

The enrichment of convergent genes in functions related to stress and their conservation in Saccharomycotina species encouraged us to question how these mRNA-mRNA interactions could confer an advantage along evolution. One hypothesis is that 3’-end RNA interactions affect mRNA access to mRNP remodeling proteins known to preferentially bind to the 3’-ends of mRNAs, such as the Pat1 and Lsm1 translational repressors comprised in P-bodies that participate in stress response (Chowdhury et al. 2007; He and Parker 2001). We analyzed CLIP data (see Materials and Methods; (Mitchell et al. 2013)) used to map the interaction sites of different P-body components, including Pat1, Lsm1, Dhh1 -that has no clear RNA sequence binding preferences- and Sbp1 that is involved in enhancing the decapping of mRNA that binds preferentially to 5’-UTR presumably resulting from its affinity to eIF4G (Rajyaguru et al. 2012; Sheth and Parker 2003; Mitchell et al. 2013).

A metagene representation of specific protein interactions has been computed for mRNA duplexes and solo mRNAs from normalized reads (in RPKM), corresponding to protein interaction sites. For solo mRNAs, as previously observed (Mitchell et al. 2013), Pat1 and Lsm1 preferentially bind the 3′-end of mRNA, Dhh1 has no positional bias and Sbp1 positions are biased towards the 5′-UTR region (fig. 6). In contrast, a significant shift in binding peaks of Pat1 and Lsm1 is observed in the 3’-UTR region of mRNA duplexes (Mann-Whitney test p-value < 0.001) (fig. 6) suggesting that 3’-RNA interactions might limit Pat1 and Lsm1 access. Spb1 also shows a decrease in the preferential binding on the 5’-UTR of mRNAs duplexes (Mann-Whitney test p-value < 0.001), which is consistent with the fact that Sbp1 binds 5’-UTR mRNA only after the binding of Pat1 and Lsm1 at their 3’-UTR, in addition to a mild decrease in their 3’-UTRs (Mann-Whitney test p-value < 0.001). The peak distribution associated to Dhh1 is not significantly different for solo and mRNA duplexes, in accordance with the fact that Dhh1 has no clear RNA sequence binding preferences (Mitchell et al., 2013). Taken together these observations argue for significant altered associations of mRNA duplexes, with Pat1 and Lsm1 leading to mRNP differing from those assembled from solo RNAs. Therefore, the fate of mRNA duplexes should differ from the fate of solo mRNAs upon stress.

**Figure 6.**
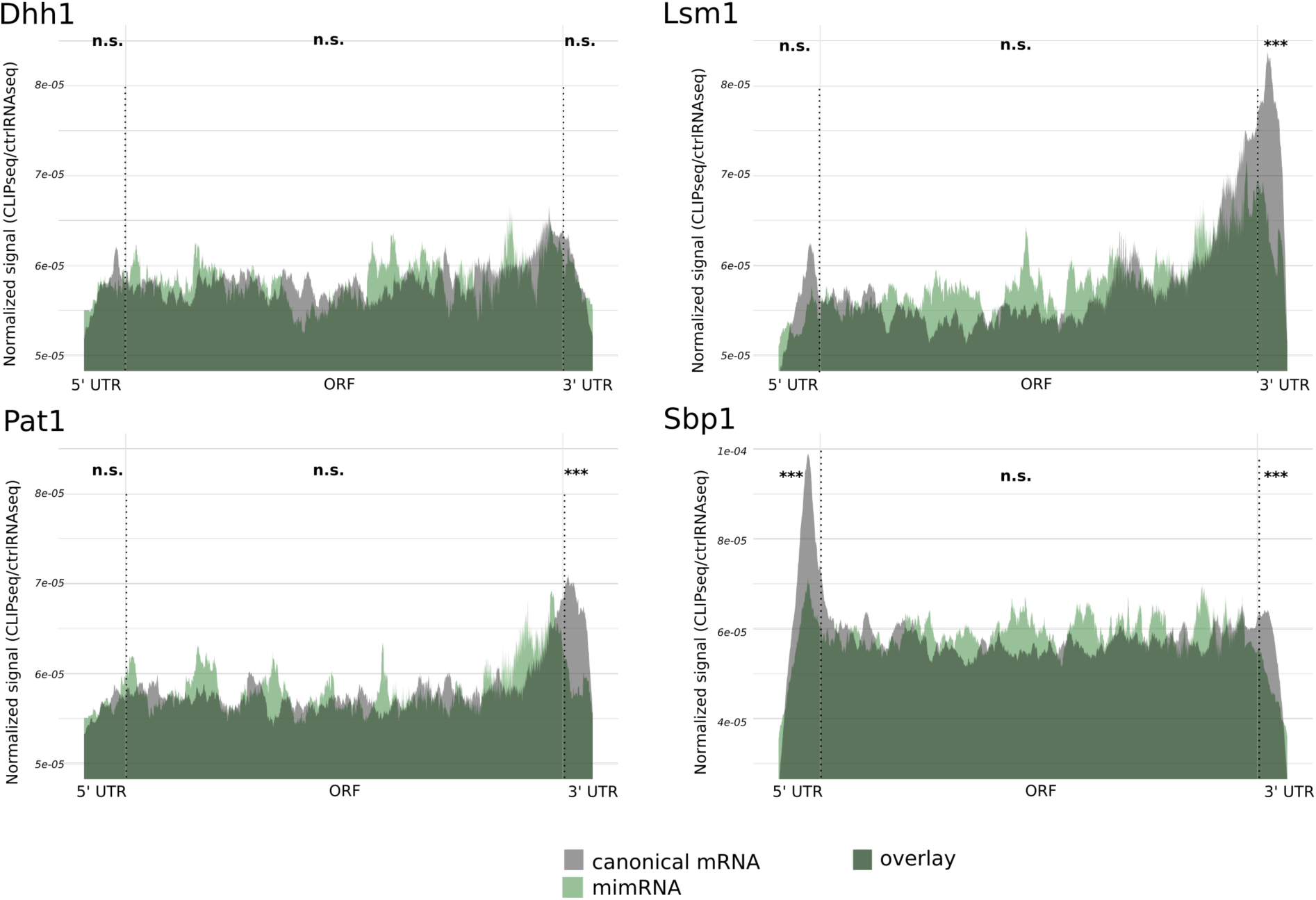
mRNA in duplexes have a marked loss of Pat1 and Lsm1 binding in their 3’-UTR. Metagene representation of sequence reads enriched using CLIP over the control sequence data for individual mRNAs. Normalized reads for solo mRNAs and mRNAs forming duplexes are represented for Pat1, Lsm1, Dhh1 and Spb1 proteins (see Materials and Methods, Mitchell et al., 2013). 5’-UTR, ORF and 3’-UTR regions are indicated. Lengths are scaled to the average 5′-UTR, ORF and 3′-UTR lengths over the entire genome. Green: mimRNAs, gray: solo mRNAs, dark green: overlay observed in regions equivalently bent within the two mRNA classes. Mann-Whitney tests p-values for comparison of the distributions between the two mRNA classes are indicated. ***: p < 1.0e-3; n.s.: p ≥ 0.05.

### The ribosome access control governed by Pat1 upon stress is limited on mRNA duplexes

In order to further examine how the decrease in interactions of Lsm1 and Pat1 on 3’-UTR regions of mRNA duplexes can affect ribosome dynamics -*i.e* translation initiation-, we took advantage of a published genome-wide analysis performed in condition of osmotic stress, during which P-bodies, involving Pat1 and Lsm1, are formed. These were used to determine ribosome mRNA associations in wild type (WT), *lsm1* and *pat1* mutant strains (Garre et al. 2018). We thus compared the ribosome accumulation at 5’-UTR of mimRNAs and solo mRNAs as identified in (Sinturel et al. 2015) in WT and *pat1* and *lsm1* mutants (supplementary table S11). The ribosome loading (*i.e*. log2 ratio of 5’ sequencing reads obtained upstream versus downstream of the mRNA translation start site (Garre et al. 2018)) in WT, *lsm1* and *pat1* mutants for each mRNA category in normal growth condition and osmotic stress is presented in fig 7. A positive shift of ribosome loading for a given mRNA between two genetic backgrounds will reflect an increase in ribosome access, thus revealing a decreased translational repression (Garre et al. 2018). A positive shift of ribosome loading in *pat1* and *lsm1* mutants was observed when all mRNAs are globally computed, as previously reported (Garre et al. 2018). We observed a similar shift for solo mRNAs and mimRNAs, which confirms the general role of Lsm1 and Pat1 in limiting ribosome access on mRNAs in normal growth conditions, (left panels, fig. 7). In stress conditions, this positive shift in ribosome loading in mutants versus WT was almost lost for mimRNAs in *pat1* mutants only (right panels, fig. 7) and also between WT and both lsm1 and pat1 strains for the 9 genes producing mimRNAs involved in stress response (supplementary table S12), but not for the 42 genes producing solos RNAs involved in stress. The probability of observing the profile of mimRNA by chance is 0.011 and of observing the profile of mimRNA involved in stress is 0.0101 (permutation tests, see Materials and Methods). The seemingly paradoxical higher translation of these 9 genes in wild type versus pat1 could result from the higher competition for translation in the mutant, in which a number of genes show de-repressed expression in stress conditions. Taken together, the above results suggest that the ribosome accumulation on mimRNAs is less dependent on the presence of Pat1 upon stress. Pat1 being considered a main translation repressor, mRNPs formed by mimRNAs would differ from mRNPs formed by solo mRNAs and be differently controlled at the translational level. We thus propose that mRNA-mRNA interactions shown to limit Pat1 mRNA binding in the 3’-region of mimRNAs (fig. 6) contribute to post-transcriptional regulations upon stress.

**Figure 7.**
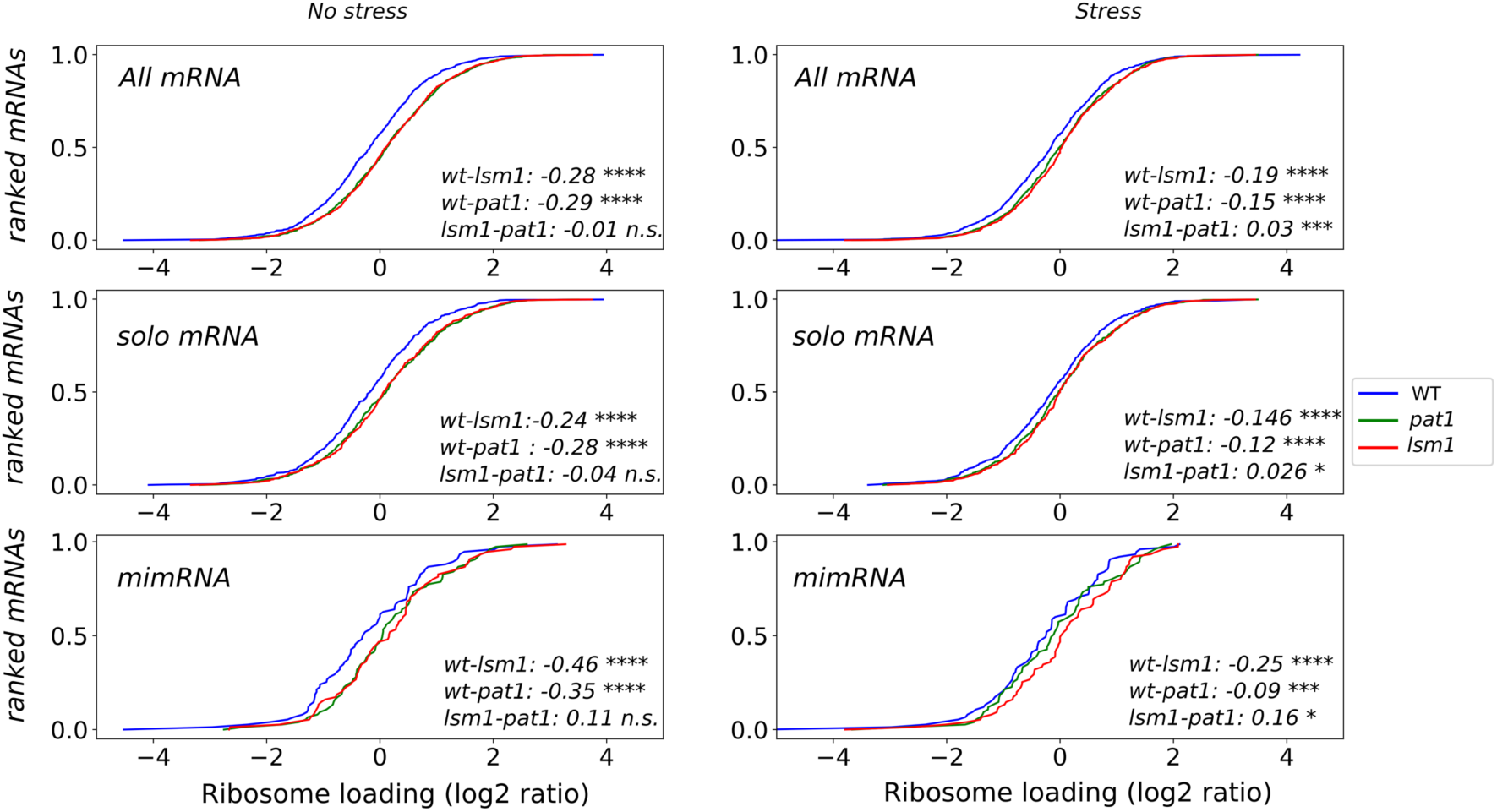
Impact of *lsm1* or *pat1* deletion on the over-accumulation of ribosomes in the 5’-UTR regions of solo mRNAs or mRNA duplexes in both control and stress conditions. Ribosome loadings are calculated as a log2 ratio between ribosome profiling reads upstream of the start codon versus downstream of the start codon for each mRNA in WT, *pat1* and *lsm1* strains, as previously determined by (Garre et al. 2018). Each mRNA is ranked along the Y axis according to its ribosome loading (X axis) calculated in different genetic backgrounds: WT (blue), *pat1* (green), *lsm1* (red). Individual panel represents ribosome loading of all mRNAs, solo mRNAs or mimRNAs in both control (no stress, left) and stress (right) conditions. For each pair of strains, the differences of the median distributions and the Wilcoxon ranked-tests p-values for comparison of the distributions are given. ****: p < 1.0e-4, ***: p < 1.0e-3; **: p < 1.0e-2; *: p < 0.05. n.s.: p ≥ 0.05.

## Discussion

In this study, we showed that convergent genes separated by short intergenic spaces are likely to produce mimRNAs that can form mRNA duplexes, independently of their 3’-UTR length. Given that the median length of intergenes separating convergent gene pairs in Saccharomycotina genomes is of 158 bp, we propose that mimRNAs can form mRNA duplexes in most of these yeasts. Indeed, intergenes between convergent pairs are the smallest ones, while those between divergent pairs are the longest ones, as previously observed among fungi (Kensche et al. 2008).

As intergenic distances is the major determinant of gene pair conservation (Poyatos and Hurst 2007), one could argue that convergent pairs, having smaller intergenic regions will inherently be more conserved, independently of selection. However, we have shown that at short intergenic distances (< 200 bp), microsynteny in convergent orientation is more conserved than in divergent and co-oriented ones. Thus, the close proximity between convergent genes can also be considered as strongly beneficial, because of tightening their linkage. A trend already observed between *Arabidopsis*, *Populus* and Rice genomes in which at distances below 250 bp there is a higher conservation of microsynteny in convergent orientation than divergent ones (Krom and Ramakrishna 2008). One could also argue that the microsynteny conservation in divergent and co-oriented orientation is counter-selected at the smallest intergenic distances that can barely contain a promoter region ranging from *ca.* 115 ± 50 bp in yeasts (Venters and Pugh 2009; Chen et al. 2011; Lubliner et al. 2013) that helps the anchoring of the transcription machinery. The bimodal distribution of intergenic distances between divergent pairs most probably reflect additional cis-regulatory constraints, as previously reported (Hermsen et al. 2008). In line with this view, among recently formed gene pairs in yeasts, divergent ones are counter-selected and are separated by very long intergenic regions (978 bp on average) (Chen et al. 2011; Sugino and Innan 2012). However, when considering the conservation of a gene relative orientation with respect to its neighbor, the same trend holds, i.e. conservation of gene orientation is higher for convergent genes than for the other orientations at small intergenic distances and we showed that the decreased conservation as the intergenic distance increases has not the same behavior for genes in convergent orientation than genes in the two other orientations. Thus, the selective pressure would be exerted on the genomic neighborhood, either conserved from a common ancestor or created by chromosomal rearrangements. Importantly, the higher conservation of convergent orientation is also observed among species that diverged after the WGD event that occurred in the yeast lineage, but to a lower extent due to the massive gene losses that occurred after the WGD, thus creating new genomic neighborhoods.

This could reflect a functional advantage of convergent genes with small intergenic spacers, related to their ability to produce mimRNAs forming mRNA duplexes and its possible influence on the post-transcriptional regulation of their expression. This is in agreement with previous analyses posing that selectively favorable co-expression appears not to be restricted to bi-directional promoters (Wang et al. 2011a). This is further supported by our observation that genes in convergent orientation present an enrichment in functions related to stress in all studied genomes. In our conservation analysis we chose not to infer relative orientations at internal nodes of the tree and therefore some branches were considered multiple times in our estimates. However, this limitation does not weaken our conclusions since they hold true when considering species pairs at similar evolutionary distances, thus reducing the bias of considering branches multiple times (supplementary fig. S2 and S3).

To investigate the structure of mRNPs produced by mRNA duplexes, we reconsidered CLIP data previously used to map the distribution of different mRNA binding proteins, Lsm1, Pat1, Dhh1, and Sbp1 on mRNAs in conditions of stress (Mitchell et al., 2013). Lsm1, Pat1, Dhh1, and Sbp1 are components of P-bodies, foci formed by stress. Interestingly, Lsm1 and Pat1 were found less frequently associated with the 3’-UTR of mimRNAs than with the 3’-UTR of others solo mRNAs. Previous analysis did not determine a particular consensus explaining why these factors bind preferentially 3’–UTR regions of solo mRNAs (Mitchell et al., 2013) but we found that 3’-end mRNA–mRNA interactions significantly counteract Lsm1 and Pat1 associations. Here we demonstrated that Dhh1-mRNA association is not affected by mRNA-mRNA interactions, confirming that Dhh1 interaction with mRNA is not region specific. Then, the less frequent associations of both factors and the moderate altered association with Sbp1 reflect a particular assembly of mRNPs. We cannot exclude that the limited Lsm1/Pat1 association also reflects a preference for other mRNA binding proteins whose access will be facilitated by the existence of double-stranded RNA sequences. In this regard, mRNP structures are complex and a multitude of other mRNA binding proteins might participate in structure assemblies of mRNA duplexes (Mitchell et al., 2013; Garre et al., 2018).

It was thus critical to assess the role of an apparent decrease in 3’-UTR associations for Lsm1 or Pat1 in the functionality of mRNA duplexes. From analysis of a genome-wide functional assay investigating the impact of Lsm1 and Pat1 on ribosome access on mRNAs (Garre et al., 2018), we found that ribosome access on mimRNAs is not modulated by Pat1, in contrast to that observed for solo mRNAs. However, we found that Lsm1 still modulates the ribosome access on mimRNAs, suggesting that Lsm1 and Pat1 have different roles for this mRNA category although their association deduced from CLIP data are similar. We thus propose that Pat1 and Lsm1 protein networks may not completely overlap and thus differently impact mRNP assemblies. In this regard, Pat1 has been proposed as a key component in promoting the formation of P-bodies (Sachdev et al. 2019). We thus propose that mimRNAs forming mRNA duplexes escape to the Pat1-dependent translation repression upon stress. The fact that stress-related genes, in general, tend to be in non-divergent relative orientation has been shown by (Wang et al. 2011b). According to these authors, stress-related genes avoid bi-promoter architecture, because they are under selection for higher noise. In our study, we provide an additional selection criterium with the demonstration that an enrichment towards stress-related processes is observed only for convergent genes forming mimRNAs, and not for genes producing solo mRNAs, whether convergent genes separated from more than 200 bp or co-oriented and divergent genes, even when they are less than 200 bp apart.

In conclusion, we showed that the conservation of the convergent orientation of genes separated by short intergenic distances is important in budding yeasts and that those convergent genes are functionally associated with stress response. Convergent genes can produce mimRNAs forming duplexes *in vivo* and our results argue for a remodeling of mRNP by those mRNA-mRNA interactions, thus providing a selective advantage for modulating gene expression upon stress, directly into the cytosol, even before any modulation of the transcription program (fig. 8). Such a post-translational regulation process is most probably conserved among budding yeasts and should be considered as a possible part of the stress response in other living cells. Thus, it should be considered as an additional factor that determines the order of coding genes.

**Figure 8.**
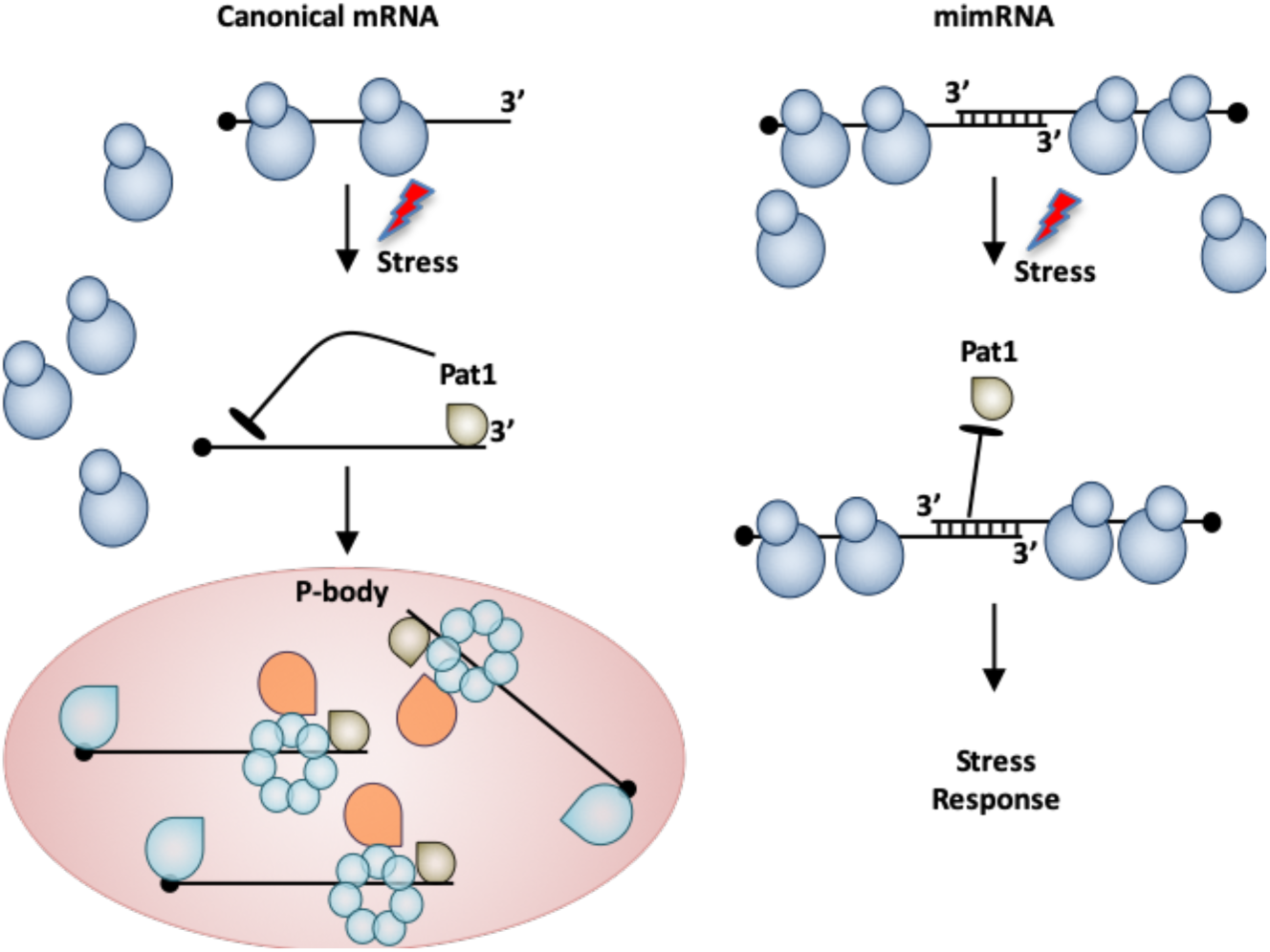
Model of the post-transcriptional regulation mediated by mRNA duplex formation. Upon stress, the translational repressor Pat1 binds preferentially to the 3’-UTR of solo mRNAs, limits ribosome access on mRNA 5’-UTRs and promotes their aggregation into P-bodies, composed by a variety of mRNA-processing factors and translational repressors. mimRNAs forming mRNA duplexes escape Pat1 repression by masking 3’-UTR access and then fully participate in stress response.

## Materials and Methods

### Data collection

The genome annotation of the *S. cerevisiae*, as well as GO Slim terms for *S. cerevisiae* genes (version 08/01/2015) were retrieved at SGD (www.yeastgenome.org), Accession numbers and address retrieval for the other 44 genomes are given in supplementary table S2.

The mimRNAs experimentally validated are those 365 mRNAs that have been sequenced with a fold coverage greater than 2 in an RNAi competent *S. cerevisiae* wild type strain (Dicer+) versus a wild type strain without Dicer (Dicer-), whereas the 248 mRNAs that have been sequenced with a fold coverage lower than one are considered as experimentally validated solo mRNAs (Sinturel et al. 2015) (supplementary table S1).

### Orthology relationships

Pairwise orthology relationships among all 45 genomes were defined between syntenic homologs retrieved with the SynCHro algorithm (Drillon et al. 2014), with the version available in June 2015. For the simulation experiment, for each species pair with N observed pairwise orthologs, we chose N pairs at random, while preserving the genome organization. The average frequencies over the 500 rounds of simulation were computed.

### Conservation of relative orientation

Between two species, we estimated the proportion of orthologs that are in the same relative orientation with respect to their gene neighbor in the two genomes. We consider the relative orientation of a gene *X_A_* in genome A and its neighboring gene *Y_A_* in genome A, compared to the relative orientation of *X_B_* in genome B, the ortholog of *X_A_* and its neighboring gene *Z_B_* in genome B. If *Y_A_* and *Z_B_* are orthologs and the relative orientation (*X_A_*, *Y_A_*) is the same as the relative orientation (*X_B_*, *Z_B_*), the relative orientation is supposed to have been present in the common ancestor, and the microsynteny between the orthologs (*X_A_*, *X_B_*) and the orthologs (*Y_A_*, *Z_B_*) is conserved. If *Y_A_* and *Z_B_* have no orthologous relationship and the relative orientation (*X_A_*, *Y_A_*) is the same as the relative orientation (*X_B_*, *Z_B_*), the relative orientation is conserved and has been acquired independently in the two genomes.

### Phylogenetic analyses

By transitivity, we inferred 224 groups of syntenic homologs composed of only one gene per species in the 45 yeasts studied. A multiple alignment of each group of orthologs was generated at the amino acid level with the MAFFT algorithm (v7.310, auto implementation, default parameters) (Katoh and Toh 2008). Concatenation of the 224 alignments was used to estimate a concatenation tree with IQtree v1.6.7 (Nguyen et al. 2015; Kalyaanamoorthy et al. 2017). The best-fit estimated model is LG+F+I+G4. Maximum likelihood distances between each species pair were estimated from the concatenated alignment and used as the evolutionary distance between species. The pairs of species were split into bins, each bin corresponding to 1/12 of the maximum evolutionary distance (1.50871) observed among all species pairs. This criterion has been chosen in order to have at least 5 species in each bin (supplementary Table S9).

### Mapping of protein RNA binding sites

Analysis are based on the CLIP sequencing datasets for the RNA binding proteins (RBP) Dhh1, Lsm1, Pat1 and Sbp1 from (Mitchell et al. 2013), downloaded at www.ncbi.nlm.nih.gov/geo/. Adapter sequences were excluded from the reads with Cutadapt v1.1 (Martin 2011) and sequences less than 22 nucleotides long were removed. Bowtie2 v2.2.3 (end-to-end mode) (Langmead and Salzberg 2012) was used to align CLIP sequencing data (22-40 bp long) against 5’-UTR, ORF and 3’-UTR from 4415 coding transcripts of the reference genome (version R57-1-1, downloaded from http://www.yeastgenome.org) and (Nagalakshmi et al. 2008) for UTR coordinates. Aligned reads received a penalty score of -6 per mismatch, -5+(-3*gap length) per gap and were excluded if penalty score was less than the default threshold (between -13.8 and -24.6 for 22 and 40 bp reads respectively). Thus, aligned reads were allowed for less than a mismatch per 10bp, (1 mismatch per 9.52 to 9.75 bp, respectively) dynamically taking into account UV-light induced mutations consecutive to the sample processing. Duplicated PCR reads, as well as reads mapping to non-coding RNAs were excluded with samtools v1.2 (Li et al. 2009) and uniquely mapped sequenced reads with a MAPQ score > 20 were kept (average MAPQ=33.26).

For each RBP-associated data, mapping of the peak interactions was constructed from a metagene aggregation procedure. The alignment depth for each gene at each nucleotide position has been determined with *samtools* (Li et al. 2009). In order to compensate control values without any sequencing signal, enrichment in the depth of sequencing signal per nucleotide coordinate *S_n_* was defined as:

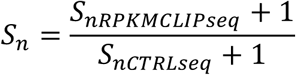

where *S_nCTRLseq_* is the sequencing signal at position *n* in the CLIP-seq experiment and *S_nCTRLseq_* is the sequencing signal at position *n* in the control experiment (RNA library without crosslinking and immunoprecipitation) of (Mitchell et al. 2013). Signal *S_n_* has been normalized to the total number of aligned reads per experiment (*i.e.* per CLIP file), and to the length of each gene (reads per kilobase of transcript per millions of aligned reads, aka. RPKM).

In order to compare the relative binding sites of each protein along the transcripts, we performed a metagene analysis. Each single gene nucleotide coordinates *n*’ have been adjusted to the longest sequence of either 5’-UTR, ORF or 3’-UTR regions to prevent any loss of information according the formula:

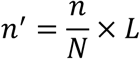

With *n* the position within the gene, *N* the gene length and *L* the longest nucleotide sequence in a defined region (5’-UTR, ORF and 3’-UTR) per CLIP experiment. Information computed for each region of a single gene were then concatenated to construct the final metagene of final length:

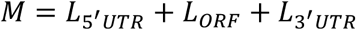

A direct interpolation was then conducted to compute the normalized depth of sequencing at each nucleotide position for the whole metagene length. Finally, the summed signal from each *S_n_*’ has been normalized according to the number of transcripts interacting with the RBP, in sort that in the final metagene representation:

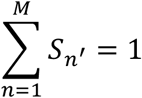

### Analysis of ribosome 5’-UTR protection

Analysis of co-translational mRNA decay by global 5’P sequencing which allows the determination of ribosome mRNA protection was previously described (Garre et al. 2018). We selected from this study only those genes having a minimum of 20 sequencing reads passing in each experiments (WT and mutant strains, in normal and stress conditions (960 genes in total, supplementary tables S11 and S12). Data were analyzed using python in-house scripts.

### Analysis of functional annotations

GO enrichment analyses among the genes forming RNA duplexes in *S. cerevisiae* with respect to the entire gene set of *S. cerevisiae* were performed with the hypergeometric test. The probability of occurrence of GO Slim terms at random in the subset of convergent genes in a given species have been computed over 1000 simulation trials. GO Slim terms for *S. cerevisiae* (go_slim_mapping.20130518.tab.gz) were retrieved at SGD’s downloads site (https://www.yeastgenome.org/search?category=download). For species others than *S. cerevisiae*, the simulations were performed by considering all genes that have orthologs in *S. cerevisiae* and that are annotated with a GO Slim term.

### Statistical analysis

Mann-Whitney U tests were performed to compare distributions of 3’-UTR length, intergenic distance, and normalized CLIP seq signal along metagenes. Wilcoxon signed-rank tests were performed to compare distributions of ribosome 5’-UTR protection. Hypergeometric tests were performed to determine the enrichment of GO Slim terms for genes forming RNA duplexes in *S. cerevisiae*, compared to the entire *S. cerevisiae* gene set. The expected probability of observing GO Slim terms for convergent genes in the other yeast genomes were performed as described in the paragraph above.

A false positive risk of a = 0.05 was chosen as a significance threshold for all tests. P-values were adjusted with the Benjamini-Hochberg false discovery rate (Benjamini and Hochberg 1995) for GO Slim enrichment and with the Holm correction in the other cases (Holm 1979). All statistical calculations were performed with R functions and with functions from the Python *scipy* module.

A permutation test for the ribosome loading analysis of solos and mimRNAs was performed by randomly picking 75 mRNAs among all mimRNAs and solo mRNAs (75+466) and counting the number of times a random profile is similar to the one observed for mimRNAs. The profile is defined by (i) a median difference between WT and pat1 lower or equal to -0.09 in stress condition and greater or equal to -0.35 in normal condition, (ii) a median difference between lsm1 and pat1 greater or equal to 0.16 in stress condition and lower or equal to 0.11 in normal condition. A similar test was performed for the ribosome loading analysis of solos and mimRNAs involved in stress biological processes. 9 mRNAs were randomly picked among all mimRNAs and solo mRNAs involved in stress (9+42) and we counted the number of times a random profile is similar to the one observed for mimRNAs. The profile is defined by a median difference between WT and pat1/lsm1 lower or equal to -0.23/0.06 in stress condition and greater or equal to 0.19/0.49 in normal condition.

## Acknowledgments

This study was supported by basic funding from CNRS and Sorbonne Université, by the ‘‘Initiative d’Excellence’’ program from the French State (Grant ‘DYNAMO’, ANR-11-LABX-0011- 01) and the AAP Emergence Sorbonne Université, SU-16-R-EMR-03. J.G. was supported by a fellowship from the Edmond de Rotschild Foundation. We thank B. Laurent for technical support, B. Billoud, M. Cavaiulo and F.A. Wollman for fruitful discussions and critical reading of the manuscript.

